# The northern gene flow into southeastern East Asians inferred from genome-wide array genotyping

**DOI:** 10.1101/2021.07.25.453681

**Authors:** Guanglin He, Yingxiang Li, Xing Zou, Hui-Yuan Yeh, Renkuan Tang, Peixin Wang, Jingya Bai, Xiaomin Yang, Zheng Wang, Jianxin Guo, Jinwen Chen, Jing Chen, Meiqing Yang, Jing Zhao, Jin Sun, Kongyang Zhu, Hao Ma, Rui Wang, Wenjiao Yang, Rong Hu, Lan-Hai Wei, Yiping Hou, Mengge Wang, Gang Chen, Chuan-Chao Wang

## Abstract

The population history of Southeast China remains poorly understood due to the sparse sampling of present-day populations and far less modeling with ancient genomic data. We here newly reported genome-wide genotyping data from 207 present-day Han Chinese and Hmong-Mien-speaking She people from Fujian and Taiwan, southeast China. We co-analyzed with 66 early-Neolithic to Iron-Age ancient Fujian and Taiwan individuals obtained from literature to explore the genetic continuity and admixture based on the genetic variations of high-resolution time transect. We found the genetic differentiation between northern and southern East Asians defined by a north-south East Asian genetic cline and the studied southern East Asians were clustered in the southern end of this cline. We also found that southeastern coastal continental modern East Asians harbored the genetic differentiation with other southern Tai-Kadai, Hmong-Mien, Austronesian and Austroasiatic speakers, as well as geographically close Neolithic-to-Iron Age populations, but relatedly close to post-Neolithic Yellow River ancients, which suggested the influence of southward gene flow on the modern southern coastal gene pool. Besides, we also identified one new Hmong-Mien genetic cline in East Asia with the coastal Fujian She localizing at the intersection position between Hmong-Mien and Han clines in the principal component analysis. She people show stronger genetic affinity with southern East Asian indigenous populations with the main ancestry deriving from Hanben-related populations. The southeastern Han Chinese could be modeled with the primary ancestry deriving from the group related to the Yellow River Basin millet farmers and the remaining from groups related to southeastern ancient indigenous rice farmers, which was consistent with the northern China origin of modern southeastern Han Chinese and in line with the historically and archaeologically attested southward migrations of Han people and their ancestors. Interestingly, *f_4_*-statistics and three-way admixture model results showed both coastal ancient sources related to Austronesian speakers and inland ancient sources related to Austroasiatic speakers complexed the modern observed fine-scale genetic structure here. Our estimated north-south admixture time ranges based on the decay of the linkage disequilibrium spanned from the Bronze age to historic periods, suggesting the recent large-scale population migrations and subsequent admixture participated in the formation of modern Han in Southeast Asia.

## INTRODUCTION

Eastern Eurasia with larger geographical regions was enriched with plentiful cultural, linguistic, and genetic diversity. More than ten different language families/groups were widely existed here, including Uralic, Altai/trans-Eurasian (Turkic, Mongolic, Tungusic, Koreanic and Japonic) in Siberia and northern China; Sino-Tibetan, Austronesian, Austroasiatic, Hmong-Mien and Tai-Kadai in East Asia and Southeast Asia. Recent genetic studies based on the modern and/or ancient genomes of East Asians have subsequently elucidated the strong association between the genetic structure, linguistic categories and geographical divisions, which were consistent with the proposed hypothesizes of the language-farming co-dispersal model (He et al., 2020d; Ning et al., 2020; Wang et al., 2020a; Yang et al., 2020; Zhang & Fu, 2020). Modern and ancient genetic studies based on the genetic variations of modern Sino-Tibetan and ancient Yellow River farmers have elucidated that the northern millet farmer expansion from northern China dispersed the Sino-Tibetan language (Wang et al., 2020a; Wang et al., 2020b), which was consistent with the common northern China origin of Sino-Tibetan languages based on the linguistic phylogeny (Sagart et al., 2019; Zhang et al., 2019). Similar studies based on the ancient DNA from southern China and Southeast Asia also found that the multiple large-scale southward dispersals of southern Chinese agriculturalists dissimilated the Austroasiatic language via the first large-scale population movements, and spread the Austronesian via coastal migration routes, Tai-Kadai, and Tibetan-Burman via the inland routes (Lipson et al., 2018; McColl et al., 2018). These gradually perfected language and genetic landscape reconstructions were benefited from the advances of ancient genome sequencing and new algorithms of genome-wide data (Patterson et al., 2012; Nielsen et al., 2017). Human population formation processes in western Eurasia have been fully explored and reconstructed, including the population substructure of Paleolithic Hunter-gatherers, cultural and genetic transition from Hunter-gathering to farming in the Neolithic revolution and the expansion of steppe pastoralists and their roles in the Indo-European language dissimilation (Mathieson et al., 2018; Olalde et al., 2018). However, the population transformations of East Asians are retained in its infancy stage. Thus, more genetic work focused on geographically/ethnically/spatiotemporally populations was needed to explore the genetic evolutionary history of East Asians.

China is of great interest in terms of deep population history and ethnolinguistic/genetic diversity, where is also the birthplace of pottery and agriculture domesticated centers of northern foxtail (*Setaria italica*) and Broomcorn (*Panicum miliaceum*) millet and Yangtze River rice (*Oryza sativa subsp. japonica*). Population genetic studies based on the low-density genetic markers, including forensic-related short tandem repeats (STRs), Insertion-Deletion (Indels), copy number variations (CNV) and single nucleotide polymorphisms (SNPs) in the autosome (Zou et al., 2018; Chen et al., 2019; He et al., 2019c; He et al., 2019a), X-chromosome (He et al., 2020a), and Y-chromosome (He et al., 2019b), have found that modern populations in China featured extensive ethnolinguistic/genetic diversity and significant genetic differentiation among populations from different language families. Spatiotemporally ancient DNA analysis results have further deconstructed that large-scale population movements and accompanying admixture could promote the formation of the increased genetic diversity and decreased genetic differentiation (Harney et al., 2018; Wang et al., 2020a). Our recent genome-wide studies focused on populations from lowland East Asia and highland Tibetan Plateau, as well as northern and southern East Asians also showed significant genetic differentiation and complex demographic processes among them (He et al., 2020b; He et al., 2020d; Wang et al., 2020c), which also emphasized the important role of modern geographically isolated or marginalized people in the demographic history reconstruction. Thus, finer-scale population genetic structure explorations from other peripheral regions of East Asia were important for the understanding of the gene pool and demographic history of ancient and modern populations.

Han Chinese are the dominant population in China and possess demographic importance for molecular anthropology, population, and forensic genetics. Wang et al. recently reconstructed one of the most important admixture models based on the ancient East Asian genomes and found that modern Han Chinese originated from the common ancestor of Sinitic and Tibetan-Burman speakers in the middle and upper Yellow River associated with the millet farmers (Wang et al., 2020a). Yang et al. also found that southward migrations of the ancient northern East Asians played an important role in the formation of the gene pool of the southern East Asians (Yang et al., 2020). Thus, comprehensive population genetic surveys of ancient and modern southern East Asians were important to elucidate the population relationships of modern southern East Asians and their northern and southern possible ancestors. Until now, 66 early-Neolithic to historic individuals from the Fujian and Taiwan have been reported (Wang et al., 2020a; Yang et al., 2020), which provided a high-resolution time transect of the peripheral region of southeast China to study the regional specific population dynamics. Here, we genotyped genome-wide data of over 700,000 SNPs in 207 unrelated individuals from 11 geographically/ethnically diverse populations (**Figure 1A**) and combined all available modern and ancient East Asians to reconstruct the population history of southeastern Asian based on the typical and advanced population genetic methods (PCA, ADMIXTURE, Fst, TreeMix, ALDER, *f-statistics* including *f_3_*, *f_4_*, *qpAdm/qpWave, qpGraph*). We found a complex population admixture history in southeastern continental East Asia since the Early Neolithic period, which is different from the observed long-term genetic continuity in other marginal regions of East Asia, such as the Tibetan Plateau (Jeong et al., 2016) and Amur River basin (Siska et al., 2017; Wang et al., 2020a).

**Figure 1.**
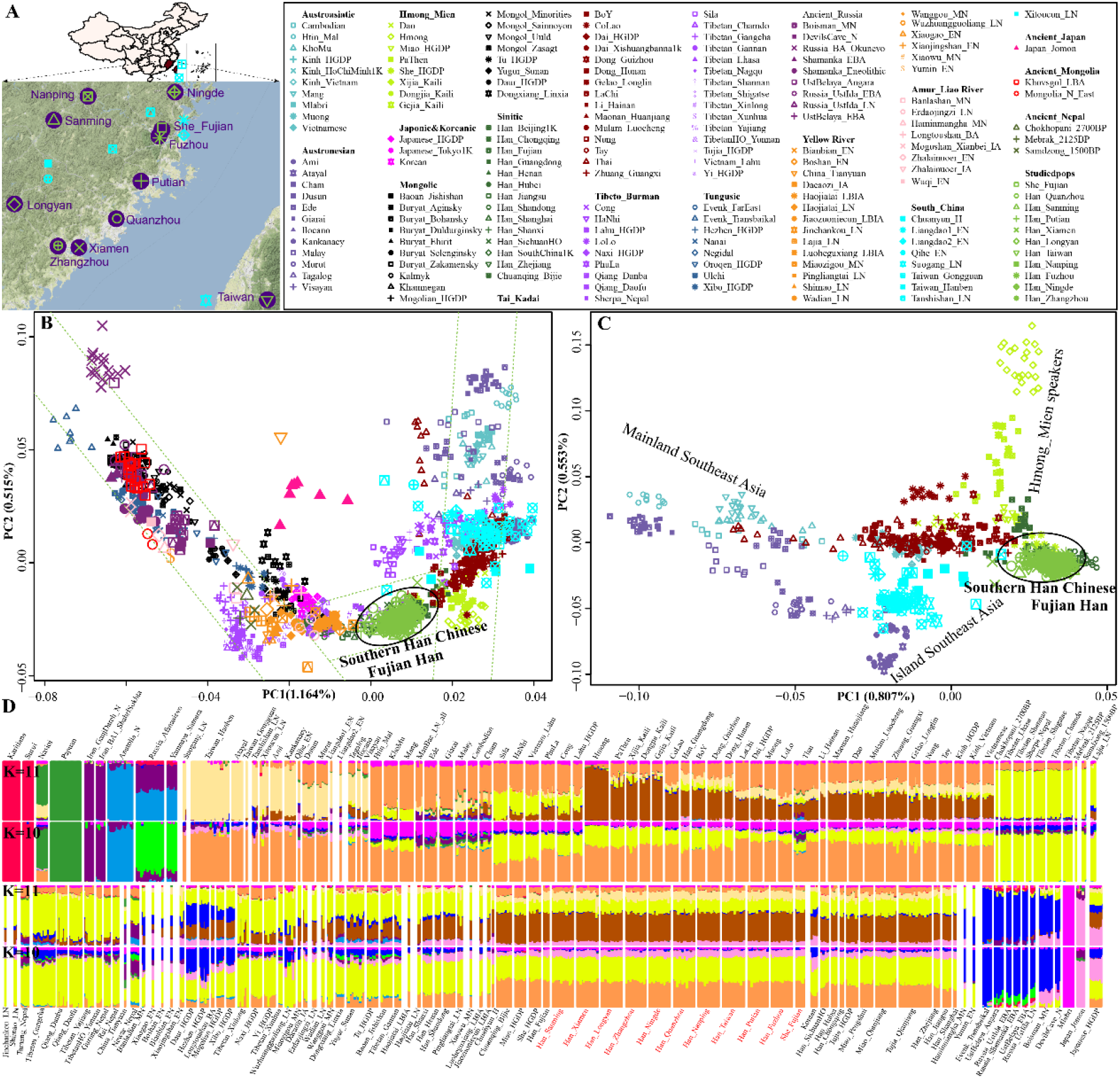
Population genetic structure inferred from the descriptive methods. (**A**) Geographical positions of 11 newly studied populations from Fujian province in Southeastern China and positions of previously published ancient Southeastern coastal East Asians. (**B~C**) The principal component analysis focused on the genetic variations between 11 studied populations and all East Asians (**B**) or southern modern and ancient East Asians (**C**). Ancient people from South Siberia, Mongolia, Nepal, China were projected onto the modern genetic background. All modern populations were categorized with their language family’s belongings. (**D**) ADMIXTURE results showed the individual/population ancestral compositions among the modern and ancient Non-Africans.

## MATERIALS AND METHODS

### Ethical approval, Sample collection and genotyping

We obtained informed consent from 207 individuals from 11 populations in southeastern Asia and collected the saliva with a saliva collection tube. Our study protocol has been approved via the medical review board at Xiamen University (Approval Number: XDYX2019009). All procedures were conducted also in accordance with the recommendations stated in the Helsinki Declaration of 2000 (Association, 2001). Genomic DNA was extracted using the PureLink Genomic DNA Mini Kit (Thermo Fisher Scientific) and qualified via Quantifiler^™^ Human DNA Quantification Kit (Applied Biosystems, Foster City, CA, USA). All samples were genotyped at the WeGene company (Shenzhen). We used the default parameters to conduct the genotype calling. We finally used Plink 1.9 to filter the new genotyped data and only remained the data with the individual genotype site missing rate less than 0.01 (mind: 0.01) and SNPs missing rate less than 0.01 (geno: 0.01). We finally obtained a dataset of 699,537 SNPs in 207 unrelated individuals after further filtered data via HWE.

### Data preparation and two working datasets

We merged newly obtained genotype data with previously published modern and ancient genomic data and obtained two working datasets of the Human-Origin-merged dataset (72K dataset) and 1240K-merged dataset (193K dataset). The 72K dataset comprised more geographically/ethnically modern populations genotyped via the Affymetrix Human-Origins chip platform, which was suitable for downstream descriptive analyses (PCA and ADMIXTURE), and the 193K dataset with higher-density SNPs suitable for following quantitative analyses (https://reich.hms.harvard.edu/downloadable-genotypes-present-day-and-ancient-dna-data-compiled-published-papers). We also included recently published ancient genome-wide data from Nepal (Jeong et al., 2016), Mongolia (Wang et al., 2020a), Siberia(Mathieson et al., 2015; Sikora et al., 2019), China (Ning et al., 2020; Wang et al., 2020a; Yang et al., 2020) and Southeast Asia (Lipson et al., 2018; McColl et al., 2018) in the population genomic analyses.

### Global ancestry analysis

#### Principal component analysis

We carried out the overall-East-Asian principal component analysis (PCA) by merging the newly genotyped Fujian and Taiwan individuals with reference panels from both northern and southern East Asians and southern-East-Asian PCA focused on the Sinitic, Austronesian, Austroasiatic, Hmong-Mien, Tai-Kadai speakers and ancient southern East Asians using the smartpca built-in EIGENSOFT packages (Patterson et al., 2012). All included ancient populations from Russia, Mongolia, Japan, China and Nepal were projected onto the modern backgrounds of the two-dimensional plots.

#### Model-based ADMIXTURE analysis

We first used Plink 1.9 (Purcell et al., 2007; Chang et al., 2015) to prune (linkage disequilibrium pruning) the successive variants with a squared correlation greater than 0.4 in 200 SNPs and sliding windows with 25 SNP in each step (--indep-pairwise 200 25 0.4). The pruned dataset consisted of new genotype data and previously published modern and ancient non-Africans, which was used to explore the East Asian substructure. We ran ADMIXTURE 1.3.0 (Alexander et al., 2009) using unsupervised mode and predefined ancestral populations ranging from 2 to 20 (K: 2~20). We calculated the cross-validation errors to choose the best-fitted model.

#### Pairwise Fst genetic distances

We used Plink 1.9 (Purcell et al., 2007; Chang et al., 2015)and our in-house script to calculate the pairwise Fst indexes (Weir & Cockerham, 1984).

#### F-statistics

We calculated admixture-*f_3_*-statistics in the form *f_3_(Source1, Source2; Fujian/Taiwan populations)* using the *qp3pop* program with default parameters in ADMIXTOOLS (Patterson et al., 2012) to explore the potential existing admixture signals and calculated *outgroup-f_3_(Eurasian1; Eurasian2; Mbuti)* to measure the shared genetic drifts between the focused populations and their reference groups. We also estimated the *f_4_*-statistics values using the *qpDstat* program and calculated the standard errors using the default block jackknife (Patterson et al., 2012).

#### Phylogenetic relationship reconstructions

We first constructed the Neighbor-joining phylogenetic tree via the TreeMix (Pickrell & Pritchard, 2012) with the migration events ranging from 2 to 20 to explore the relationship between Fujian/Taiwan populations and their neighboring modern and ancient reference populations. And then we also constructed another Neighbor-joining tree based on the inverse outgroup-*f_3_*-based genetic distance matrix.

#### Multidimensional scaling analysis

We carried out multidimensional scaling analysis based on the inverse outgroup-*f_3_*-based genetic distance matrix using SPSS software and then visualized in R software.

#### Streams of ancestry and inference of mixture proportions

We used *qpWave* as implemented in ADMIXTOOLS (Patterson et al., 2012) to explore the minimum number of ancestral sources needed to illuminate the observed genetic variations in newly studied populations and used *qpAdm* to quantify the ancestral proportions of each ancestral populations. All eleven newly-studied populations were used as the left or tested populations and nine worldwide populations (Mbuti, Russia_Ust_Ishim, Russia_Kostenki14, Papuan, Australian, Mixe, Russia_MA1_HG, Onge, Atayal) were used as the right/reference populations. The additional parameter of ‘allsnps: YES’ was used here.

#### Admixture graph modeling

We used *qpGraph* (Patterson et al., 2012) to construct two admixture graphs based on the one simple seed skeletal framework and one complex framework with two archaic introgression events (Neanderthal introgression to non-African and Denisovan introgression to Australasian (Green et al., 2010; Reich et al., 2010)). We finally fitted all studied eleven populations on these two models to explore the ancestral admixture proportions for their northern and southern East Asian ancestral proxies.

#### Dating of gene-flow events

We used multiple Admixture-induced Linkage Disequilibrium for Evolutionary Relationships (MALDER) to estimate the admixture times with 31 potential ancestral candidates (Loh et al., 2013). We used 29 years of the length of one generation and calculated the years before the Common Era (BCE): Y=1950-29*(ALDER-calculated years-1).

#### Y-chromosomal and mtDNA haplogroup assignment

We used our in-house scripts and the identified derived upstream alleles and ancestral downstream alleles to assign to haplogroups.

## RESULTS

### Population genetic structure of southeastern modern and ancient East Asians

We reported approximately 700K genome-wide SNPs in 207 Han Chinese and She individuals from Fujian and Taiwan in southeastern China and merged with 66 geographically close ancient people, neighboring East Asians to study the population dynamics of the southeastern coast in East Asia. We first conducted overall-East-Asian PCA based on the 72K dataset to assess the genetic affinity between newly genotyped populations and published ancient and present-day populations. We found three genetic clines including northern cline (Tungusic and Mongolic speakers at one end and Tibeto-Burman speakers at the other end), southern cline (Austronesian and Austroasiatic speakers at one end and Hmong-Mien at the other end) and the intermediate Han Chinese cline between Tibeto-Burman and Tai-Kadai people (**Figure 1B**). Fujian and Taiwan populations were localized at the southmost end of the Han Chinese cline and showed a close relationship with modern Tai-Kadai-speaking populations and ancient southern East Asians (Liangdao, Hanben, Tanshishan, Xitoucun and so on). Regional-southern-East-Asian PCA focused on Han and southern East Asians showed a clear finer-scale population substructure. Here, we found four genetic sub-clines, including Austronesian, Austroasiatic, Sinitic and Hmong-Mien clines, which were consistent with the linguistic affiliations. The Tai-Kadai-speaking populations and ancient southern East Asians were positioned in the middle of the above four gradient groups (**Figure 1C**). Interestingly, we identified one Hmong-Mien genetic cline here which showed a close genetic relationship with newly studied Han Chinese, Hmong-Mien-speaking She and Tai-Kadai-speaking populations.

To further explore the ancestry composition of Fujian and Taiwan populations and their genetic similarity with their geographically close ancient populations, we carried out model-based ADMIXTURE analysis among 4,130 non-African modern and ancient individuals from 355 populations (mainly from eastern Eurasia and also included representative populations from Europe, Oceania and America). We found obvious genetic differences between modern and ancient southeastern Asians but a relatively stronger extent of genetic homogenization between modern and ancient northern East Asians although genetic differentiation could be identified. When K=10, we observed three ancestry components in newly studied populations: Orange ancestry predominated in southeastern modern Austronesian (Ami and Atayal) and ancient groups (Hanben, Tanshishan and Xitoucun), Yellow ancestry enriched in Nepal ancients and modern Tibetans and pink ancestry maximized in ancient Jomon people. When K=11, a new bronze ancestry was maximized in Hmong-Mien-speaking Hmong, which was also enriched in Pathen and dominant in our studied She and Han Chinese groups.

### Admixture signatures and shared genetic drifts

The genetic affinity among southeastern East Asians was further confirmed via the smallest pairwise Fst genetic distances (**Table S1**) and the largest shared genetic drift within them (**Table S2**). Fujian and Taiwan populations had the smaller genetic distances with geographically close present-day and ancient populations (from Fujian Tanshishan and Xitoucun and Taiwan Hanben sites) and also with northern Sinitic speakers. Outgroup-*f_3_*-statistics also showed that Fujian and Taiwan populations shared the most genetic drift with all East Asian Sinitic speakers, southern Tai-Kadai and Austronesian speakers, and Late Neolithic to Iron Age southern East Asians (*f_3_*-values reached ~0.3, **Figure 2A~B**). We followingly performed the multidimensional scaling analysis (MDS) among modern and ancient East Asians based on the inverse *f_3_*-based genetic distances (1/*f_3_*) and found similar patterns of genetic clusters as in PCA and ADMIXTURE results. Ancient coastal southern East Asians from Fujian and Taiwan clustered together and localized close with Austronesian populations, while ancient coastal and inland northern East Asians grouped separately and showed close relationships with modern and ancient populations from Tibet Plateau and northern China. Han Chinese and She populations in Fujian and Taiwan clustered tightly together between northern and southern East Asians or between northern Sinitic and southern Tai-Kadai speaking populations on a finer-scale (**Figure 2C**). We performed admixture-*f_3_(Source1, source2; eleven Fujian/Taiwan populations)* to explore the potential ancestral sources among 354 modern and ancient Eurasians with statistically significant *f_3_*-values with Z-scores less than-3. For Hmong-Mien-speaking She, we observed the mixed signatures in 311 out of 66410 ancestral candidates’ pairs with one source usually from modern or ancient southern East Asians and the other from northern Tibetan-Burman/Tungusic/Mongolic-related populations (**Figure 2D~G**), such as the most negative *f_3_* value was *f_3_* (Ami, Chamdo Tibetan; Fujian She) = −6.423*SE.

**Figure 2.**
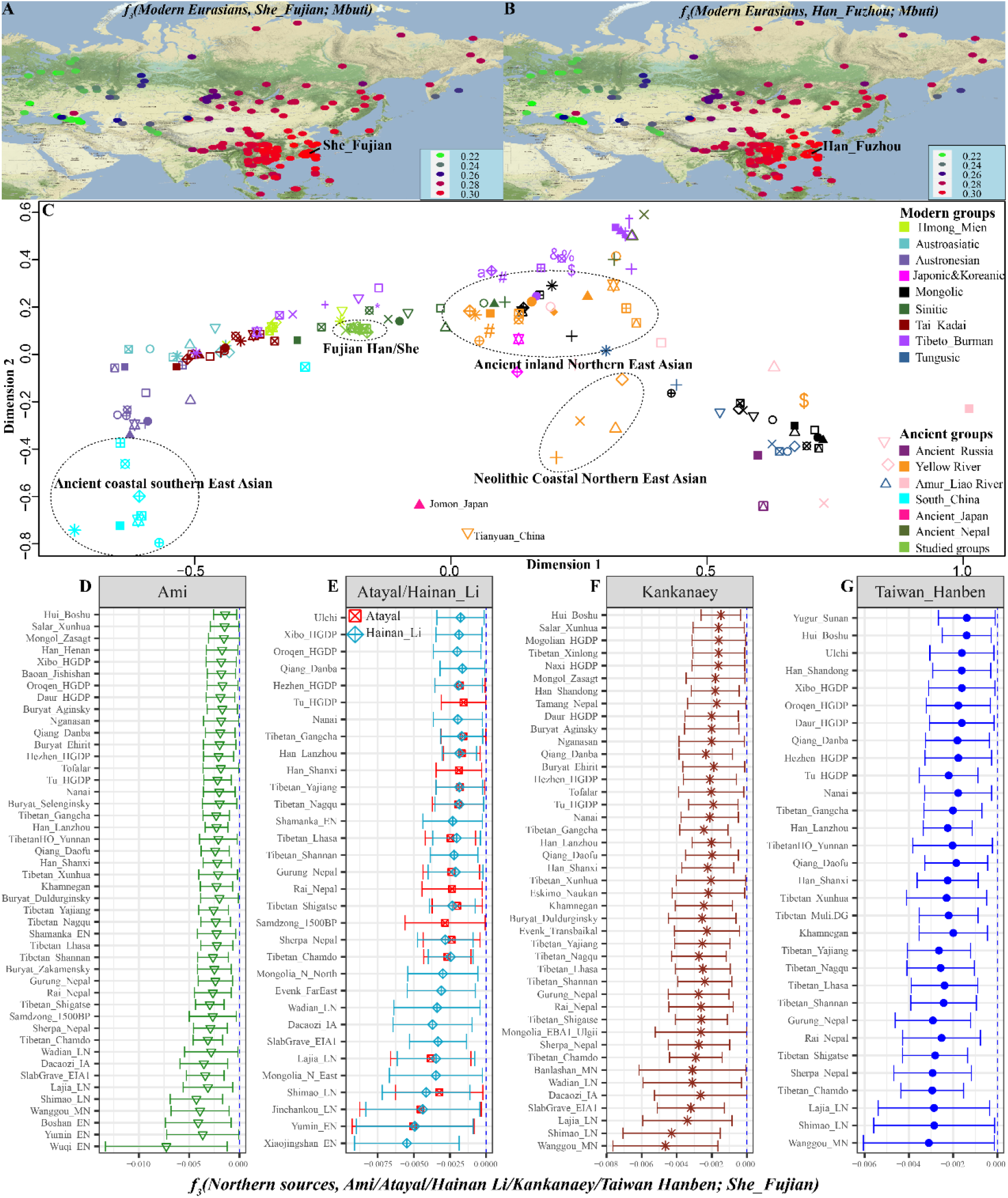
Shared genetic drifts and mixture signals were revealed via outgroup-*f_3_*-statistics and admixture-*f_3_*-statistics. (**A~B**) Heatmap of *f_3_*-statistic values showed the genetic affinity between Fujian She (**A**) or Fuzhou Han (**B**) and other Eurasian modern reference populations. (**C**) Multidimensional scaling analysis based on the matrix of the inverse shared genetic distance (1/outgroup-*f_3_*) showed the genetic relationship between 11 Fujian populations and other modern and ancient East Asians. (**D~G**) Statistically significant admixture-*f_3_*-statistic signatures focused on Fujian She when we used the southern East Asians of Ami. Atayal, Li, Kankanaey and ancient Taiwan Hanben as the plausible southern sources.

### Genetic continuity and admixture

Qualitative analysis of PCA and ADMIXTURE model-based clustering showed the genetic differentiation between northern and southern East Asians defined by a north-south cline and intermediate positions of our newly reported Han Chinese and She populations in the north-south cline. Qualitative testing based on the admixture-*f_3_* and outgroup-*f_3_*-statistics further showed the closer genetic affinity of our newly reported populations with southern East Asians and Bronze to Iron Age northern East Asians. To examine genetic continuity and potential admixture events between southeastern coastal Han Chinese and She with ancient East Asians, we performed five different types of *f_4_*-statistics. Firstly, we conducted *f_4_(Han/She1; Han/She2; Eurasians, Mbuti)* and found no population substructure among 10 Han Chinese populations (**Table S3**). However, we detected more allele sharing between southern indigenous populations (Dong Kankanaey) or northern coastal Neolithic Xiaojingshan samples with She people compared with Han Chinese (**Figure 3A**).

**Figure 3.**
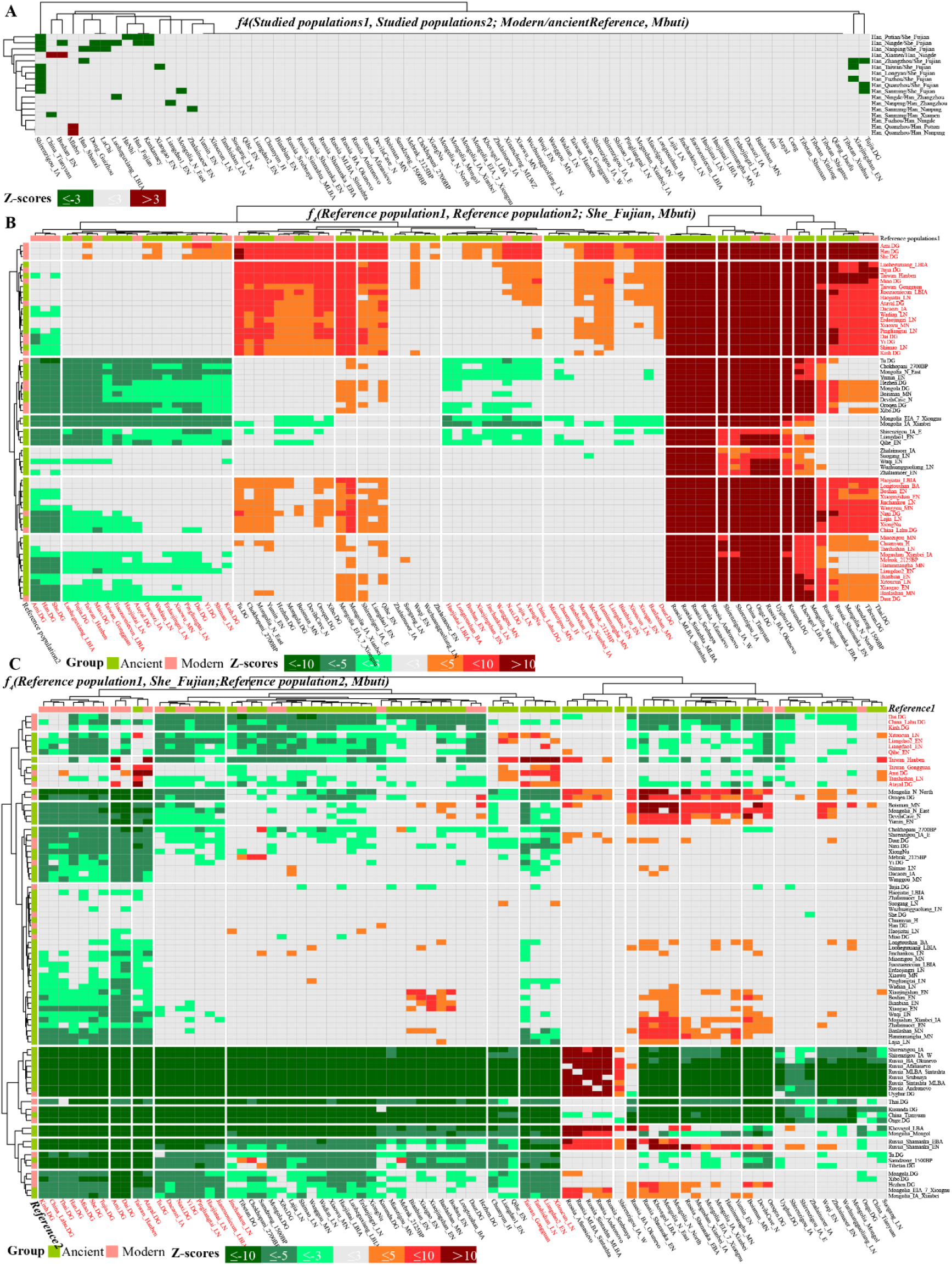
Four-population-based statistics showed the asymmetrical genetic affinity between Fujian populations and their corresponding references. (**A**) Symmetrical-*f_4_*-statistics in the form *f_4_(Studied Fujian population1, Studied Fujian population2; modern and ancient Eurasian reference populations, Mbuti)*. (**B**) Affinity-*f_4_*-statistics in the form *f_4_(Reference population1, Reference population2; Fujian She, Mbuti)* shared the stronger genetic affinity between Fujian She and modern and ancient southern East Asians. (**C**) Asymmetrical *f_4_*-statistics in the form *f_4_(Reference populations, Fujian She; Ancestral sources or Reference populations, Mbuti)* showed the genetic continuity and admixture between modern Fujian populations and their ancient predecessors.

Secondly, to further explore the genetic heterogeneity among Sino-Tibetan speakers, Chinese ancient populations and southern Chinese and Southeast Asians from four language families and validate the genetic homogeneity among the newly-studied southeastern Chinese populations, we performed pairwise *qpWave*-based analysis among 113 populations used in regional-PCA analysis based on the powerful set of outgroups (Mbuti, Ust_Ishim, Kostenki14, Papuan, Australian, Mixe, MA1, Onge, Mongolia_N_East, Wuqi_EN, Bianbian_EN, Liangdao2_EN, Jomon, Yumin_EN, Chokhopani and Yamnaya). The *qpWave* analysis was conducted via various sets of *f_4_*-statistics in the form *f_4_(left_1_,left_i_;right_1_,right_i_)* to explore the gene flow events from the used right populations to the left populations (Patterson et al., 2012). We found genetic differences between linguistically different populations and genetic differentiation between modern southeastern East Asians and their geographically close predecessors (**Table S4**), but confirmed the genomic similarity within the linguistically/geographically close populations, especially in our studied populations (**Figure 4**). Here, we could identify pairwise p_rank0 larger than 0.05, which suggested the genetic cladality existed between two used left populations compared with the used right outgroups. We could identify five clade clusters, including four homogeneous clusters (northern homogeneous group, southern homogeneous group, northern ancient homogeneous group and southern Hmong-Mien/Tai-Kadai homogenous group). All studied Fujian and Taiwan populations showed similar homogeneous signals of genetic cladality with each other, as well as with previously reported Fujian Han, Bijie Chuanqing, Miao and She included in the Human Genome Diversity Project. Interestingly, homogeneous signals between the newly - studied populations and ancient northern populations (Jinchankou, Dacaozi, Pingliangtai, Wadian, Haojiatai and Xiaowu) from the Central plain, where is the birthplace of Chinese Civilization, suggesting the southward gene flow events and their significant impact on the gene pool composition of modern southeastern coastal populations. Here, we also observed genetic heterogeneity between modern southeastern Han populations and geographically close Neolithic predecessors, inland modern Tai-Kadai/Hmong-Mien speakers, as well as with northern inland ancient populations from the upper Yellow River, suggested their differentiated demographic admixture history.

**Figure 4.**
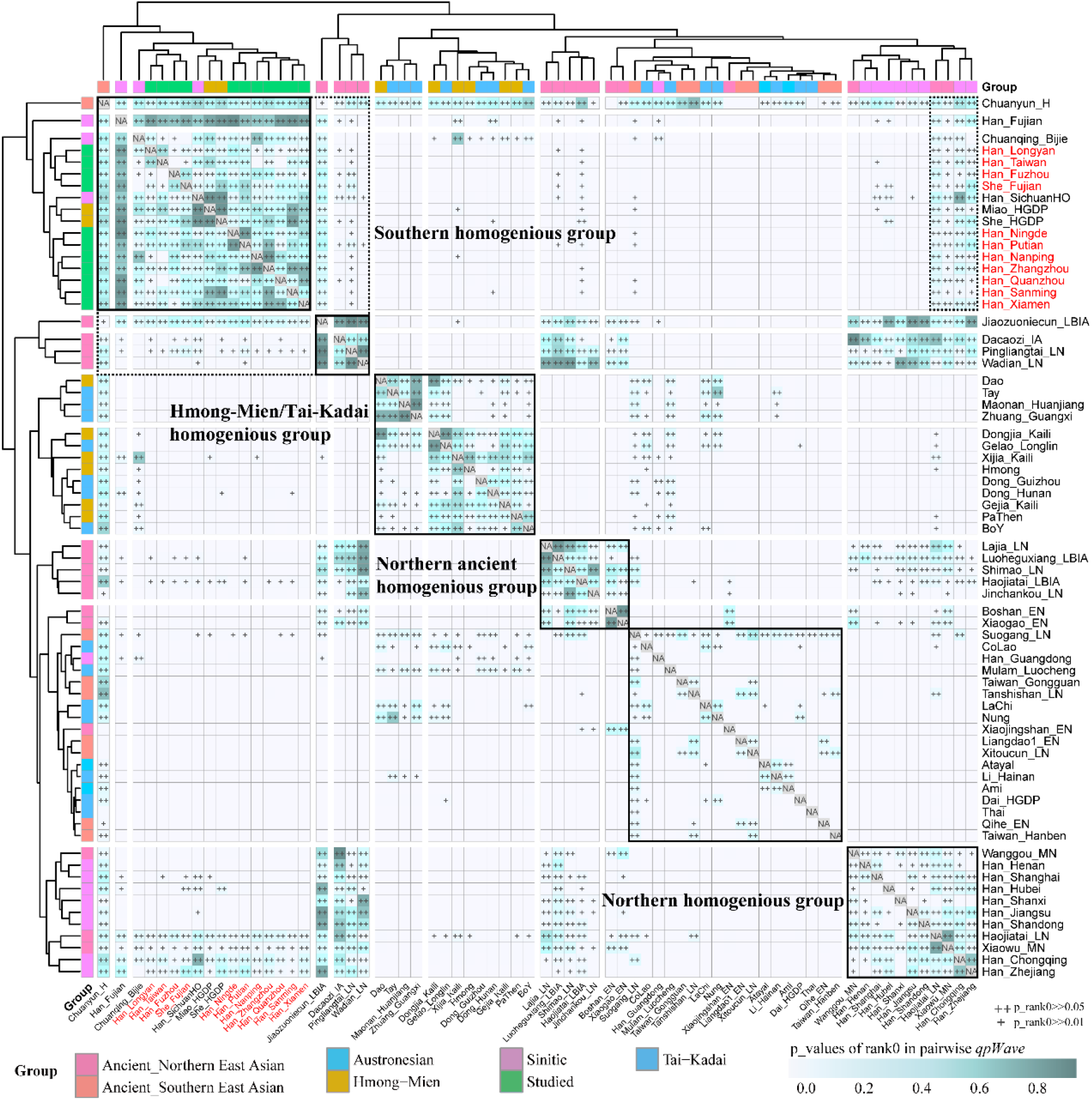
Pairwise p-values of *qpWave* for each pair of populations. Values greater than 0.05 are marked with two parallel plus (++) signs and greater than 0.01 are marked with one plus sign.

Thirdly, to further explore the genetic affinity formally or additional gene-flow events from surrounding heterogeneous ancestral source candidates among the 193K dataset, we performed *f_4_(Eastern Eurasians1, Eastern Eurasians2; 11 studied populations, Mbuti)* and found similar patterns of shared ancestry among Han/She and their reference populations (**Table S5**). No significant negative *f_4_*-statistic values were observed in *f_4_(Eastern Eurasians1, Eastern Eurasians2; Fujian She, Mbuti)* when southern East Asians (modern Ami, Han, She, Tujia, Miao and Atayal and ancient Hanben and Gongguan) and northern middle Neolithic to Iron Age (Luoheguxiang_LBIA, Jiaozuoniecun_LBIA, Haojiatai_LN, Erdaojingzi_LN and Xiaowu_MN) used as the first reference populations (Eastern Eurasians1, the right populations listed in **Figure 3B**), which suggested significant more allele sharing of Fujian She with them than with other reference populations (Eastern Eurasians2, the bottom populations listed in **Figure 3B**). Compared with Neolithic people from Siberia and Inner Mongolia and Early Neolithic Liangdao people in southeast China, Fujian She possessed excess sharing alleles with Late Neolithic to modern East Asians. In detail, significant positive values in *f_4_(Northern Yellow River ancient Yangshao/Houli populations, other reference populations; southeastern Han/She, Mbuti)* observed here implied the southward dispersal of gene flow from the Yellow River into the southeast coast since the Late Neolithic (**Figure 3B**).

Fourth, we used *f_4_(reference population 1, southeastern Han/She; reference population 2, Mbuti)* to evaluate whether modern southeastern East Asians were consistent with directly descending from ancient populations from Yellow River Basin or southeastern coastal regions (here we used as reference population 2, bottom populations listed in **Figure 3C**). We identified the most negative *f_4_*-values when we used modern southern groups (Kinh, Lahu, Thai, Han, Miao, She, Tujia, Ami, Dai, Atayal) and ancient Iron Age Hanben samples as the reference population 2 (**Table S6**). We also observed the consistent genetic affinity of negative *f_4_*-values when northern modern (Tu, Naxi and Yi) and ancient populations (Dacaozi_IA, Pingliangtai_LN, Haojiatai_LN and Jiaozuoniecun_LBIA) and southern ancient coastal populations (Chuanyun_H, Liangdao1_EN, Qihe_EN and Taiwan_Gongguan, Tanshishan_LN, Liangdao2_EN and Xitoucun_LN) as the reference population 2 (**Figure 3C**). These observed negative signals showed their affinity with southeastern East Asians, consistent with the assumptions that reference population 2 was predefined as their ancient ancestral source candidates.

Fifth, to further explore whether one or more ancestral populations (mainly focused on geographically close southeastern coastal ancient East Asians, and homogeneous ancient populations from the Yellow River basin) could be fitted as the unique ancestral populations for our studied modern populations, we performed *f_4_(candidate source, 11 studied populations; reference populations, Mbuti)*. We used the population that can give a negative *f_4_*-value as “reference population 2” in the above *f_4_* statistics as the candidate source (**Table S6**). Genetic cladality between Southeastern East Asians and Wuzhuangguoliang, Suogang and Chuanyun ancient populations were observed, which may be used as direct evidence for studied populations directly descended from these three populations or related populations.

Here, it is should be cautious that all identified ancestral sources may be biased by the low-overlapping SNPs and high contamination of these samples (Wang et al., 2020a; Yang et al., 2020). In fact, we observed additional admixture signals via symmetrical *f_4_*-statistics. The populations from northern and southern East Asia (bottom population lists in **Figure 3C**) shared excess alleles with our studied Fujian populations compared to the candidate source populations (reference populations listed in the right list in **Figure 3C**). Consistent with the negative admixture signals observed in admixture-*f_3_*-statistics in the form *f_3_(candidate source, reference populations; 11 studied propulsions)*, we concluded that southeastern modern Han and She cannot be modeled as descending directly from geographically close Neolithic to historic ancient populations without additional admixture related to ancient Yellow River Basin agriculturalists. Modern southeastern East Asian was formed via complex admixed processes.

Summarily, serval lines of evidence supported the southward gene flow from northern East Asia into the southeastern continental East Asia region, which is also confirmed via *f_4_*-statistics based on the merged Human Origin dataset (**Tables S7~8**). First, a recent ancient genomic study conducted via Yang et al. demonstrated that southward migrations from the Yellow River basin significantly influenced the gene pool of Fujian Late Neolithic populations (Yang et al., 2020). As they observed that the Late Neolithic Tanshishan and Xitoucun people harbored more Yellow farmer-related ancestry compared with early Liangdao people (Yang et al., 2020), suggested by the observed negative values in *f_4_(Liangdao, Tanshishan_LN/Xitoucun_LN; Yellow River millet farmer; Mbuti)*. Second, combined analysis among ancient genomes reported via Yang et al. (Yang et al., 2020), Ning at al. (Ning et al., 2020) and Wang et al. (Wang et al., 2020a) also revealed the continuous southward from Yellow River millet further into Taiwan Iron Age Hanben and Gongguan populations in the southeastern continental East Asia marginal region, as the negative *f_4_-values* in *f_4_(Liangdao_EN/Tanshishan_LN/Xitoucun_LN, Hanben_Taiwan/Gongguan_Taiwan; ancient northern East Asian, Mbuti)*. Third, Further evidence from *f_4_*-statistics (**Figure 3**) demonstrated that present-day southeastern East Asians harbored more northern East Asian related ancestry compared to their early ancestor as evidenced via the *f_4_*-statistics in the form *f_4_(ancient southeastern continental East Asians, modern Fujian/Taiwan populations; ancient northern East Asians, Mbuti)*. Thus, millet agricultural development and millet farmer southward expansion continued to have an impact on the formation of the gene pool of the southern East Asians. These north-south admixture processes were further confirmed via the *qpWave/qpAdm*, *qpGraph* and MALDER results in the next section.

### Admixture processes and their potential phylogenetic relationship

Results from the descriptive and qualitative analyses suggested genetic variations or allele frequency of studied southeastern populations tended to be at an intermediate position between northern and southern East Asians. Thus, we further used *qpWave* analysis to study the minimum ancestral populations that could be used to explain our observed genetic pattern and found that two ancestral candidates can be best-fitted (p_rank1 > 0.05) with one powerful set of outgroups. We employed *qpWave* and *qpGraph* to estimate the ancestral proportion of their potential ancestral sources. First, we used all available northern millet farmer-related populations as the northern sources and Iron Age Hanben from Taiwan as the southern sources and used the *qpAdm* to estimate the mixing proportions. We observed She and southeastern Han derived more northern ancestry from the ancient Yellow River farming populations in the majority of the models except for the Yumin_EN-Hanben model (**Figure 5**). Second, we well-fitted two *qpGraph*-based phylogenies and added all eleven studied populations on the skeletal formworks. We found all included populations could be fitted as admixtures between groups related to the Yellow River millet farmers and Yangtze River ghost populations (**Figure 6**). Third, we have found the genetic differences between southern coastal East Asians and southern inland East Asians. Thus, to further explore the gene flow influence from inland southern East Asians related to Austroasiatic speakers or their ancestors on the genetic structure of modern coastal populations, we conducted three-way admixture models, in which twenty-one ancient northern East Asians were used as the northern sources, three ancient populations from mainland Southeast Asia were used as inland southern inland source and Ami from Taiwan as the coastal source. We could successfully model eleven studied populations as the admixture results of major ancestry from northern China, approximately equal ancestry from two southern sources (**Table S9**).

**Figure 5.**
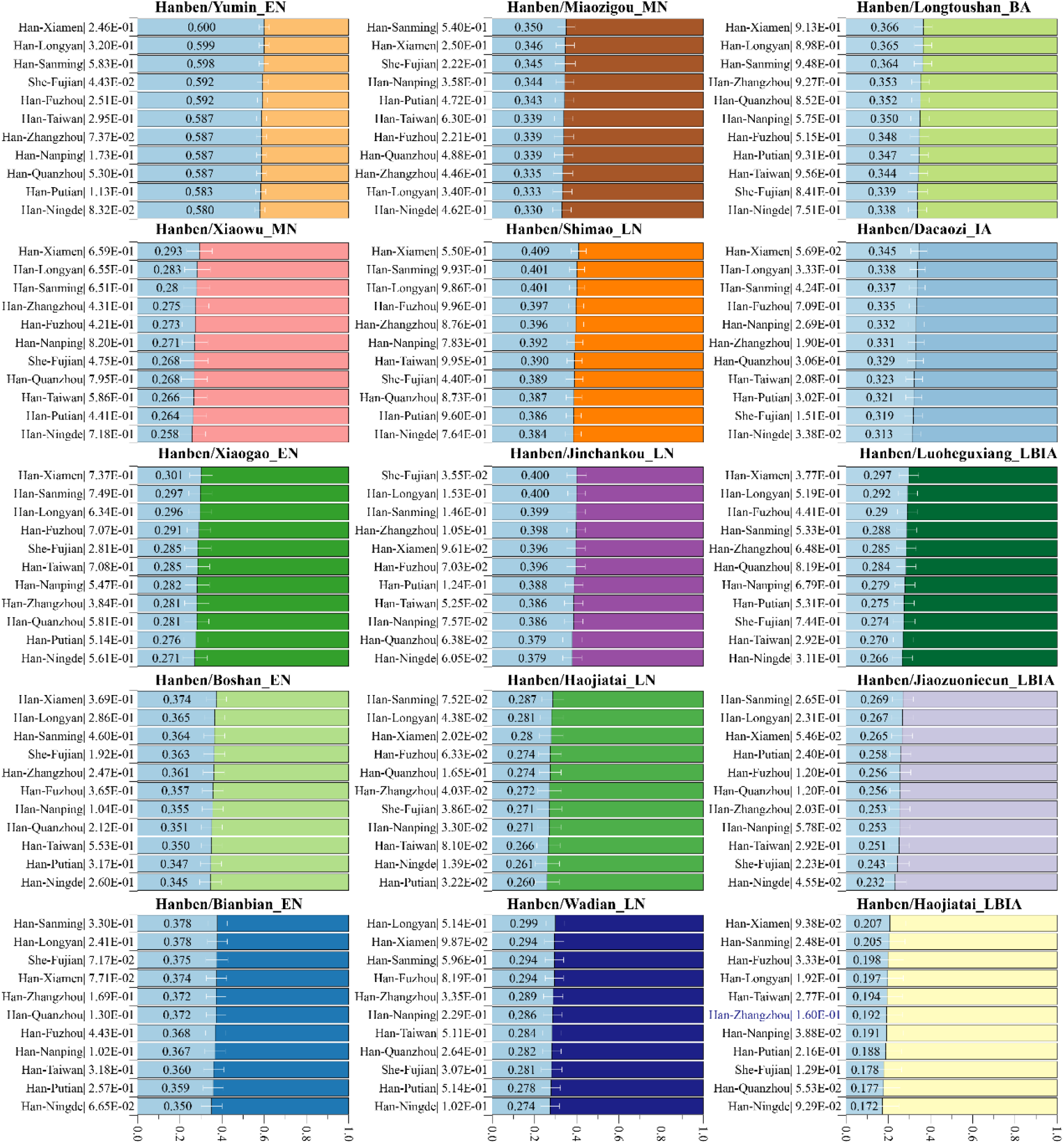
*qpAdm/qpWave* results showed the detailed ancestral composition in the best-fitted two-way admixture model. Here, we used Iron Age Taiwan Hanben as the southern ancestral sources which possessed a stronger affinity with modern Austronesian and Tai-Kadai-speaking populations. Early Neolithic to Iron Age northern East Asian from the Yellow River Basin and their surrounding regions were used as the possible northern Sources. New studied populations and their corresponding p_rank1 in the *qpWave* tests were listed in the left part of the bar plots. White err bar denoted as the standard error.

**Figure 6.**
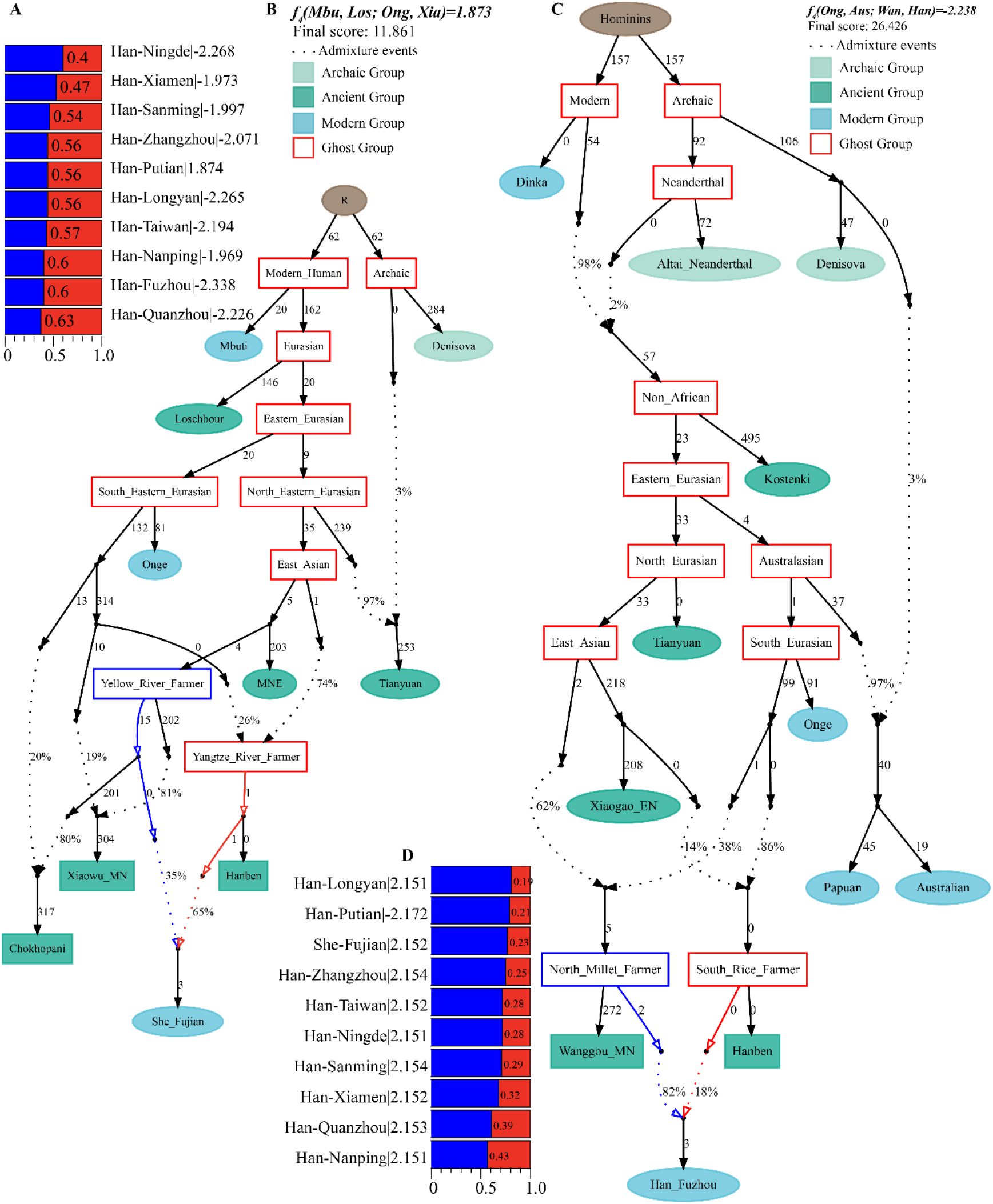
Genetic models fitted using qpGraph showed the formation of modern Southern East Asian Fujian populations. We fitted the two admixture graphs by sequentially adding populations to the seed models, and finally fitted all new studied populations in simple seed graph model (**A~B**) and complex seed graph model (**C~D**).

Furthermore, Neighbor-joining phylogenetic relationships constructed based on the matrix of 1/*f_3_* also found a tight cluster of southeastern coastal modern Han and She, which localized between northern meta-groups (Tungusic/Mongolic clade, ancient northern East Asian clade, ancient and modern Tibetan-Burman clade and northern Sinitic clade) and southern-meta groups (Hmong-Mien clade, southern Tibeto-Burman clade, Austronesian/ancient southern coastal East Asian clade, and Austroasiatic/Tai-Kadai clade). Here, we also found historic Chuanyun people clustered with Xiamen Han and Fujian Han, suggesting their more shared ancestral history (**Figure 7**). Similar patterns of phylogenetic relationships were evidenced by Fst genetic distance matrix. ALDER-based admixture times with different ancestral populations further showed the north-south population interaction and admixture occurred at least from Neolithic time and large-scale population contact occurred in the Bronze Age to the historic time (**Figure 8**).

**Figure 7.**
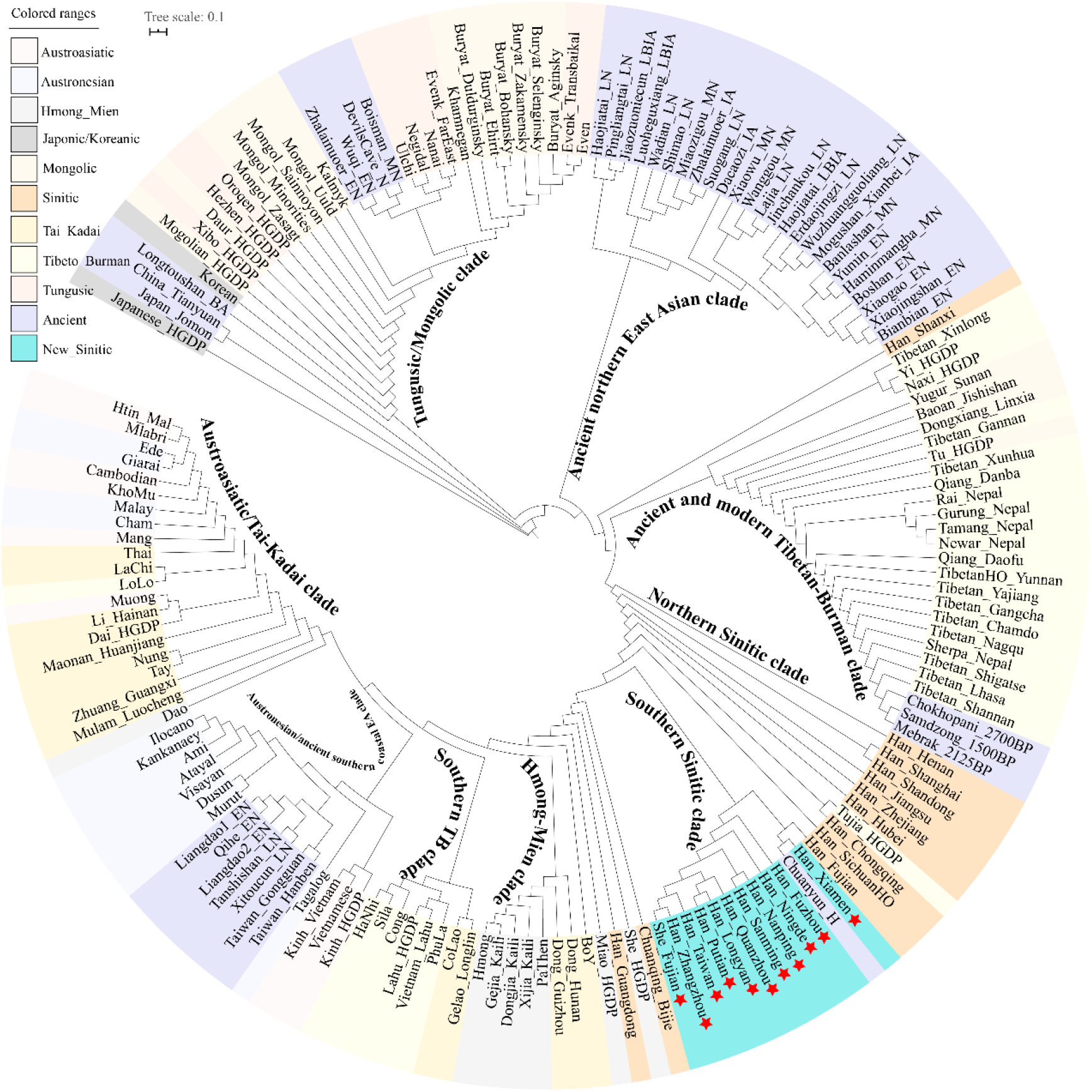
Phylogenetic relationships between eleven Fujian populations and other modern and ancient East Asian reference populations based on the shared *f_3_*-based genetic distance matrix (1/*f_3_*). All included modern populations were grouped with their linguistic affiliations.

**Figure 8.**
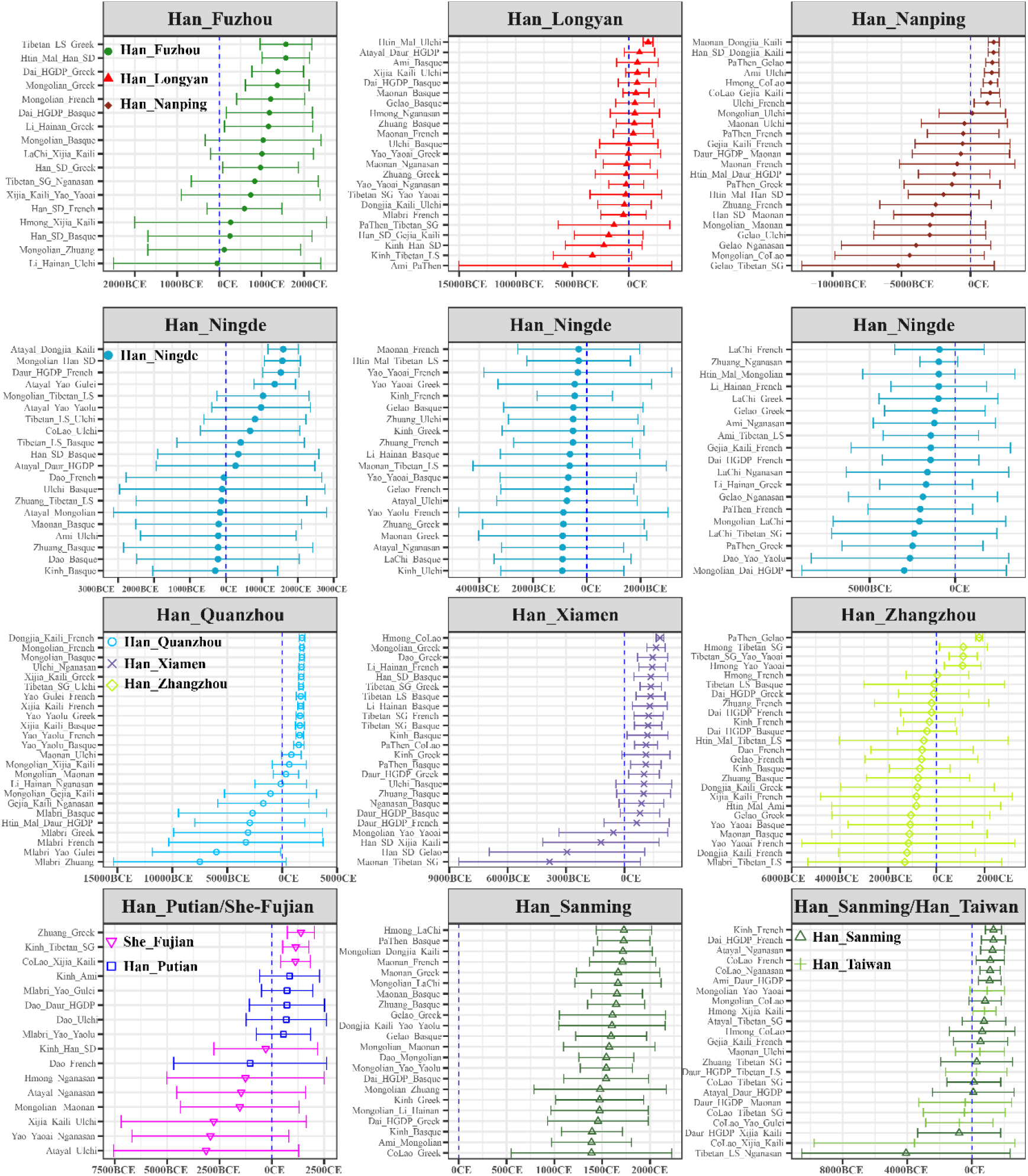
The estimated admixture times for Fujian populations based on the decay rates of admixture-induced linkage disequilibrium. All listed two-way admixture models were well-fitted models with the expected p values and Z-scores. The marked admixture years were calculated as: Y=1950-29*(calculated used generations-1).

### Uniparental genetic structure of southeastern Asians

We assigned 207 mitochondrial genomes into 118 different terminal haplogroups with the corresponding frequency ranging from 0.0048 (1) to 0.0386 (F1a1, 8). Haplogroups F1a1, D4a, M7c1c, B4a, F1a and M8a were the dominant maternal lineages in southeastern East Asians (**Table S10**). We assigned 108 Y-chromosome genomes into 66 terminal haplogroups with the haplogroup frequency ranging from 0.0093 (1) to 0.0463 (O2a2b1a1a5a2, 5). O1a was regarded as the common founding lineage of modern southern Han, Tai-Kadai and Austronesian speaking populations, which comprised 0.2315 (25/108) of studied Han and She populations. Han-dominant O2a was the major lineage consisting of 0.4722 (51/108) of studied populations. It was interesting to find that one western Eurasian specific R1a1a1b2a2b1b and one Siberian-dominant Q1a1a1a1a~ were observed in southeastern Han Chinese, suggesting the long-range of mobility and admixture in eastern Eurasia.

## DISCUSSION

Population genetic studies focused on East Asians have illustrated the geographically isolated populations or populations from the peripheral regions have a relatively homogenized structure with long-term genetic continuity. He et al. studied the genetic structure of the Tai-Kadai-speaking Li population from Hainan island using genome-wide data and found that the Li people were uniquely homogeneous with no obvious recent gene flow from mainland China (He et al., 2020b). Results of genome-wide studies focused on highland-adaptive populations in the Qinghai-Tibet Plateau also found the primary ancestry of modern Tibetans derived from the millet agriculturalists from the middle and upper Yellow River Basin in northern China (Wang et al., 2020a). This work also demonstrated that modern core Tibetans and ancient Nepal people (Samdzong, Chokhopani, and Mebrak) acquired little additional gene flow when the human population permanently occupied the highland at least from late Neolithic time (approximately 3600 years ago) (Jeong et al., 2016; Wang et al., 2020a). Genetic evidence from Tungusic speakers residing in the peripheral region of Northeast Asia and geographically close ancient DevilsCave people also illuminated their long-term genetic stability at least from 7700 years ago (Siska et al., 2017). However, populations from other regions that may act as the homeland or corridors of population movement have complex demographical changes (Chen et al., 2019; He et al., 2020d). Here we comprehensively investigated the population genetic structure and admixture history in Fujian and Taiwan at the southeastern edge of East Asia. Our genetic analysis focusing on the genetic history from early Neolithic to modern populations demonstrated the past of the southeast coast was dynamic with multiple population mixing, for example, the gene flow from northern farming populations into the southeast coast. Generally, the population history of southern East Asia was complexed via the mixing process of different gene pools observed in this study, including southward dispersal northern East Asian gene pool, indigenous southern differentiated gene pools associated with the Austronesian, Hmong-Mien, Tai-Kadai and Austronesian speakers. Our quantitative analyses of ADMIXTURE and PCA consistently demonstrated that modern southern East Asians were differentiated into inland and coastal lineages, and studied southern continental East Asians harbored gene flow from a northern lineage related to the shared lineage of all Sino-Tibetan speakers and two southern indigenous lineages. Quantitative analyses via *f*-statistics, pairwise *qpWave* and *qpAdm* further confirmed the population migrations related to these three lineages that participated in the formation of southeastern coastal modern population genetic legacy. We also identified genetic differentiation between modern southeastern continental populations, and neighboring Neolithic populations and modern Austronesian speakers suggested that the southward gene flow events shifted the ancient genetic structure, which is consistent with the geographical feature of the corridor in ancient coastal population migrations and the main starting point of the southward dispersal of Austronesian speakers and their language (Yang et al., 2020).

The origin of the Sino-Tibetan people remains heavily debated due to three different hypotheses been evidenced via linguistic, historic findings to some extent, including the North Indian origin hypothesis, Southwest China/Tibetan-Yi Corridor hypothesis, and northern China origin hypothesis. Our results also supported the primary ancestry of modern southeastern coastal Han Chinese originated from northern China, which is consistent with previous genome-wide analysis from southernmost, Central and northern modern Han Chinese (He et al., 2020c; He et al., 2020b; He et al., 2020d). Linguistic evidence has also illuminated the common origination of Sinitic and Tibeto-Burman languages (Sagart et al., 2019; Zhang et al., 2019) from north China, in accordance with the archeologically documented Yangshao/Majiayao-related culture dispersal. Our results from the *qpAdm* results have found the main ancestry of southeastern Han deriving from northern millet farmer related populations, which were further confirmed via the *qpGraph*-based demographic models. Further, we identified the finer-scale population substructure among southern East Asians consistent with the linguistic affiliation, but differentiated with the observed genetic structure of southeastern coastal populations. Except for the previous identified Austronesian and Austroasiatic mainly from the Island and Mainland of Southeast Asia, here, we identified one new Hmong-Mien-speaking population genetic cline consisting of Hmong, Pathen and Dao from the mainland of southeastern Asia (Liu et al., 2020), Gejia, Dongjia, Xijia (Sun et al., 2020), Miao and She from southern China. Stronger southern East Asian affinity of Fujian She and its close relationship with populations from inland Hmong-Mien-speaking population genetic cline, consistent with the linguistic classification of She people. Here, we note we should pay greater attention to the limitation of sampling bias of Hmong-Mien She, which should be further confirmed with more ancient DNA from southern China and denser sampling of modern Hmong-Mien-speaking populations.

## CONCLUSION

Genome-wide evidence from modern and ancient southeastern coastal East Asians demonstrated the continuous southward gene flow since the early Holocene has shaped their genetic landscape. Present-day southeastern Han and Hmong-Mien She derived ancestry from at least two ancient populations respectively associated with Yellow River millet farmers and Yangtze River rice agriculturalists. The obtained primary ancestry and the reconstructed phylogeny have provided genome-wide evidence of northern China origin of southeastern Han Chinese and ALDER-based admixture times further illuminated the large-scale southward migrations that occurred during Bronze Age and historic times. Finer-scale population substructure within southern East Asians visualized here could provide significant clues for further deep populations history reconstruction, rethink the language-farming-co-dispersal hypothesis and even present some genetic reference for experimental design of cohort studies related to precision medicine. More ancient DNA studies and denser modern population sampling should be further conducted to explore more details of the demographic developments of modern southern East Asians.

## ACKNOWLEDGEMENTS

This study was supported by the National Natural Science Foundation of China (31801040), the China Postdoctoral Science Foundation (2021M691879 and 2021M691882), Nanqiang Outstanding Young Talents Program of Xiamen University (X2123302), and Fundamental Research Funds for the Central Universities (ZK1144). YXL was also supported by National Postdoctoral Program for Innovative Talents (BX20180180).

## DATA AVAILABILITY

Following the regulations and informed consent of this project and the regulations of the Human Genetic Resources Administration of China (HGRAC), the obtained raw data can be shared via personal communication with corresponding authors. We make the data available upon request by asking the person requesting the data to agree in writing to the following restrictions: 1, the data can be only used for studying population history; 2, the data cannot be used for commercial purposes; 3, the data cannot be used to identify the sample donors; 4, the data cannot be used for studying natural/cultural selections, medical or other related studies.

## Legends of supplementary Tables

**Table S1**. The pairwise Fst genetic distance among 49 East Asian modern and ancient populations.

**Table S2**. The shared genetic drift between eleven southeastern continental Han/She populations and other ancient/modern Eurasians.

**Table S3**. Differentiated shared alleles among eleven studied populations compared to modern and ancient Eurasians estimated via symmetrical f4-statistics in the form f4(Han/She1, Han/She2; eleven studied populations, Mbuti) based on the merged Human Origin dataset.

**Table S4**. Pairwise p_rank0 values in *qpWave* analysis among 113 East Asian modern and ancient populations.

**Table S5**. Symmetrical f4-statistics in the form f4(Reference population1, reference population2; eleven studied populations, Mbuti) to test their genetic affinity between Eurasian reference populations based on the merged 1240K dataset.

**Table S6**. Affinity-f4-statistics in the form f4(Reference population1, eleven studied populations; reference population2, Mbuti) to explore the genetic continuity and admixture between potential ancestral sources and Fujian/Taiwan populations based on the merged 1240K dataset.

**Table S7**. Symmetrical f4-statistics in the form f4(Reference population1, reference population2; eleven studied populations, Mbuti) to test their genetic affinity between Eurasian reference populations based on the merged Human Origin dataset.

**Table S8**. Affinity-f4-statistics in the form f4(Reference population1, eleven studied populations; reference population2, Mbuti) to explore the genetic continuity and admixture between potential ancestral sources and Fujian/Taiwan populations based on the merged Human Origin dataset.

**Table S9**. qpAdm-based ancestral admixture proportion in eleven studied populations via three-way admixture models.

**Table S10**. Haplogroup distribution in eleven studied populations.

## REFERENCE

Alexander DH, Novembre J, Lange K. 2009. Fast model-based estimation of ancestry in unrelated individuals. Genome Res 19: 1655–1664.

Association WM. 2001. World medical association declaration of helsinki. Ethical principles for medical research involving human subjects. Bulletin of the World Health Organization 79: 373.

Chang CC, Chow CC, Tellier LC, Vattikuti S, Purcell SM, Lee JJ. 2015. Second-generation plink: Rising to the challenge of larger and richer datasets. Gigascience 4: 7.

Chen P, Wu J, Luo L, Gao H, Wang M, Zou X, Li Y, Chen G, Luo H, Yu L, Han Y, Jia F, He G. 2019. Population genetic analysis of modern and ancient DNA variations yields new insights into the formation, genetic structure, and phylogenetic relationship of northern han chinese. Front Genet 10: 1045.

Green RE, Krause J, Briggs AW, Maricic T, Stenzel U, Kircher M, Patterson N, Li H, Zhai W, Fritz MH, Hansen NF, Durand EY, Malaspinas AS, Jensen JD, Marques-Bonet T, Alkan C, Prufer K, Meyer M, Burbano HA, Good JM, Schultz R, Aximu-Petri A, Butthof A, Hober B, Hoffner B, Siegemund M, Weihmann A, Nusbaum C, Lander ES, Russ C, Novod N, Affourtit J, Egholm M, Verna C, Rudan P, Brajkovic D, Kucan Z, Gusic I, Doronichev VB, Golovanova LV, Lalueza-Fox C, de la Rasilla M, Fortea J, Rosas A, Schmitz RW, Johnson PLF, Eichler EE, Falush D, Birney E, Mullikin JC, Slatkin M, Nielsen R, Kelso J, Lachmann M, Reich D, Paabo S. 2010. A draft sequence of the neandertal genome. Science 328: 710–722.

Harney E, May H, Shalem D, Rohland N, Mallick S, Lazaridis I, Sarig R, Stewardson K, Nordenfelt S, Patterson N, Hershkovitz I, Reich D. 2018. Ancient DNA from chalcolithic israel reveals the role of population mixture in cultural transformation. Nat Commun 9: 3336.

He G, Zou X, Wang M, Liu J, Wang F, Hou Y, Wang Z. 2020a. Population genetics, diversity, forensic characteristics of four chinese populations inferred from x-chromosomal short tandem repeats. Leg Med (Tokyo) 43: 101677.

He G, Wang Z, Zou X, Wang M, Liu J, Wang S, Ye Z, Chen P, Hou Y. 2019a. Tai-kadai-speaking gelao population: Forensic features, genetic diversity and population structure. Forensic Sci Int Genet 40: e231–e239.

He G, Wang Z, Su Y, Zou X, Wang M, Chen X, Gao B, Liu J, Wang S, Hou Y. 2019b. Genetic structure and forensic characteristics of tibeto-burman-speaking u-tsang and kham tibetan highlanders revealed by 27 y-chromosomal strs. Sci Rep 9: 7739.

He G, Ren Z, Guo J, Zhang F, Zou X, Zhang H, Wang Q, Ji J, Yang M, Zhang Z, Zhang J, Nabijiang Y, Huang J, Wang CC. 2019c. Population genetics, diversity and forensic characteristics of tai-kadai-speaking bouyei revealed by insertion/deletions markers. Mol Genet Genomics 294: 1343–1357.

He G, Wang Z, Guo J, Wang M, Zou X, Tang R, Liu J, Zhang H, Li Y, Hu R, Wei LH, Chen G, Wang CC, Hou Y. 2020b. Inferring the population history of tai-kadai-speaking people and southernmost han chinese on hainan island by genome-wide array genotyping. Eur J Hum Genet 28: 1111–1123.

He G, Wang M, Li Y, Zou X, Yeh HY, Tang R, Yang X, Wang Z, Guo J, Luo T, Zhao J, Sun J, Hu R, Wei LH, Chen G, Hou Y, Wang CC. 2020c. Fine-scale north-to-south genetic admixture profile in shaanxi han chinese revealed by genome-wide demographic history reconstruction. Journal of Systematics and Evolution: 0-.

He GL, Li YX, Wang MG, Zou X, Yeh HY, Yang XM, Wang Z, Tang RK, Zhu SM, Guo JX, Luo T, Zhao J, Sun J, Xia ZY, Fan HL, Hu R, Wei LH, Chen G, Hou YP, Wang CC. 2020d. Fine-scale genetic structure of tujia and central han chinese revealing massive genetic admixture under language borrowing. Journal of Systematics and Evolution n/a.

Jeong C, Ozga AT, Witonsky DB, Malmstrom H, Edlund H, Hofman CA, Hagan RW, Jakobsson M, Lewis CM, Aldenderfer MS, Di Rienzo A, Warinner C. 2016. Long-term genetic stability and a high-altitude east asian origin for the peoples of the high valleys of the himalayan arc. Proc Natl Acad Sci U S A 113: 7485–7490.

Lipson M, Cheronet O, Mallick S, Rohland N, Oxenham M, Pietrusewsky M, Pryce TO, Willis A, Matsumura H, Buckley H, Domett K, Nguyen GH, Trinh HH, Kyaw AA, Win TT, Pradier B, Broomandkhoshbacht N, Candilio F, Changmai P, Fernandes D, Ferry M, Gamarra B, Harney E, Kampuansai J, Kutanan W, Michel M, Novak M, Oppenheimer J, Sirak K, Stewardson K, Zhang Z, Flegontov P, Pinhasi R, Reich D. 2018. Ancient genomes document multiple waves of migration in southeast asian prehistory. Science 361: 92–95.

Liu D, Duong NT, Ton ND, Van Phong N, Pakendorf B, Van Hai N, Stoneking M. 2020. Extensive ethnolinguistic diversity in vietnam reflects multiple sources of genetic diversity. Mol Biol Evol 37: 2503–2519.

Loh PR, Lipson M, Patterson N, Moorjani P, Pickrell JK, Reich D, Berger B. 2013. Inferring admixture histories of human populations using linkage disequilibrium. Genetics 193: 1233–1254.

Mathieson I, Lazaridis I, Rohland N, Mallick S, Patterson N, Roodenberg SA, Harney E, Stewardson K, Fernandes D, Novak M, Sirak K, Gamba C, Jones ER, Llamas B, Dryomov S, Pickrell J, Arsuaga JL, de Castro JM, Carbonell E, Gerritsen F, Khokhlov A, Kuznetsov P, Lozano M, Meller H, Mochalov O, Moiseyev V, Guerra MA, Roodenberg J, Verges JM, Krause J, Cooper A, Alt KW, Brown D, Anthony D, Lalueza-Fox C, Haak W, Pinhasi R, Reich D. 2015. Genome-wide patterns of selection in 230 ancient eurasians. Nature 528: 499–503.

Mathieson I, Alpaslan-Roodenberg S, Posth C, Szecsenyi-Nagy A, Rohland N, Mallick S, Olalde I, Broomandkhoshbacht N, Candilio F, Cheronet O, Fernandes D, Ferry M, Gamarra B, Fortes GG, Haak W, Harney E, Jones E, Keating D, Krause-Kyora B, Kucukkalipci I, Michel M, Mittnik A, Nagele K, Novak M, Oppenheimer J, Patterson N, Pfrengle S, Sirak K, Stewardson K, Vai S, Alexandrov S, Alt KW, Andreescu R, Antonovic D, Ash A, Atanassova N, Bacvarov K, Gusztav MB, Bocherens H, Bolus M, Boroneant A, Boyadzhiev Y, Budnik A, Burmaz J, Chohadzhiev S, Conard NJ, Cottiaux R, Cuka M, Cupillard C, Drucker DG, Elenski N, Francken M, Galabova B, Ganetsovski G, Gely B, Hajdu T, Handzhyiska V, Harvati K, Higham T, Iliev S, Jankovic I, Karavanic I, Kennett DJ, Komso D, Kozak A, Labuda D, Lari M, Lazar C, Leppek M, Leshtakov K, Vetro DL, Los D, Lozanov I, Malina M, Martini F, McSweeney K, Meller H, Mendusic M, Mirea P, Moiseyev V, Petrova V, Price TD, Simalcsik A, Sineo L, Slaus M, Slavchev V, Stanev P, Starovic A, Szeniczey T, Talamo S, Teschler-Nicola M, Thevenet C, Valchev I, Valentin F, Vasilyev S, Veljanovska F, Venelinova S, Veselovskaya E, Viola B, Virag C, Zaninovic J, Zauner S, Stockhammer PW, Catalano G, Krauss R, Caramelli D, Zarina G, Gaydarska B, Lillie M, Nikitin AG, Potekhina I, Papathanasiou A, Boric D, Bonsall C, Krause J, Pinhasi R, Reich D. 2018. The genomic history of southeastern europe. Nature 555: 197–203.

McColl H, Racimo F, Vinner L, Demeter F, Gakuhari T, Moreno-Mayar JV, van Driem G, Gram Wilken U, Seguin-Orlando A, de la Fuente Castro C, Wasef S, Shoocongdej R, Souksavatdy V, Sayavongkhamdy T, Saidin MM, Allentoft ME, Sato T, Malaspinas AS, Aghakhanian FA, Korneliussen T, Prohaska A, Margaryan A, de Barros Damgaard P, Kaewsutthi S, Lertrit P, Nguyen TMH, Hung HC, Minh Tran T, Nghia Truong H, Nguyen GH, Shahidan S, Wiradnyana K, Matsumae H, Shigehara N, Yoneda M, Ishida H, Masuyama T, Yamada Y, Tajima A, Shibata H, Toyoda A, Hanihara T, Nakagome S, Deviese T, Bacon AM, Duringer P, Ponche JL, Shackelford L, Patole-Edoumba E, Nguyen AT, Bellina-Pryce B, Galipaud JC, Kinaston R, Buckley H, Pottier C, Rasmussen S, Higham T, Foley RA, Lahr MM, Orlando L, Sikora M, Phipps ME, Oota H, Higham C, Lambert DM, Willerslev E. 2018. The prehistoric peopling of southeast asia. Science 361: 88–92.

Nielsen R, Akey JM, Jakobsson M, Pritchard JK, Tishkoff S, Willerslev E. 2017. Tracing the peopling of the world through genomics. Nature 541: 302–310.

Ning C, Li T, Wang K, Zhang F, Li T, Wu X, Gao S, Zhang Q, Zhang H, Hudson MJ, Dong G, Wu S, Fang Y, Liu C, Feng C, Li W, Han T, Li R, Wei J, Zhu Y, Zhou Y, Wang CC, Fan S, Xiong Z, Sun Z, Ye M, Sun L, Wu X, Liang F, Cao Y, Wei X, Zhu H, Zhou H, Krause J, Robbeets M, Jeong C, Cui Y. 2020. Ancient genomes from northern china suggest links between subsistence changes and human migration. Nat Commun 11: 2700.

Olalde I, Brace S, Allentoft ME, Armit I, Kristiansen K, Booth T, Rohland N, Mallick S, Szecsenyi-Nagy A, Mittnik A, Altena E, Lipson M, Lazaridis I, Harper TK, Patterson N, Broomandkhoshbacht N, Diekmann Y, Faltyskova Z, Fernandes D, Ferry M, Harney E, de Knijff P, Michel M, Oppenheimer J, Stewardson K, Barclay A, Alt KW, Liesau C, Rios P, Blasco C, Miguel JV, Garcia RM, Fernandez AA, Banffy E, Bernabo-Brea M, Billoin D, Bonsall C, Bonsall L, Allen T, Buster L, Carver S, Navarro LC, Craig OE, Cook GT, Cunliffe B, Denaire A, Dinwiddy KE, Dodwell N, Ernee M, Evans C, Kucharik M, Farre JF, Fowler C, Gazenbeek M, Pena RG, Haber-Uriarte M, Haduch E, Hey G, Jowett N, Knowles T, Massy K, Pfrengle S, Lefranc P, Lemercier O, Lefebvre A, Martinez CH, Olmo VG, Ramirez AB, Maurandi JL, Majo T, McKinley JI, McSweeney K, Mende BG, Modi A, Kulcsar G, Kiss V, Czene A, Patay R, Endrodi A, Kohler K, Hajdu T, Szeniczey T, Dani J, Bernert Z, Hoole M, Cheronet O, Keating D, Veleminsky P, Dobes M, Candilio F, Brown F, Fernandez RF, Herrero-Corral AM, Tusa S, Carnieri E, Lentini L, Valenti A, Zanini A, Waddington C, Delibes G, Guerra-Doce E, Neil B, Brittain M, Luke M, Mortimer R, Desideri J, Besse M, Brucken G, Furmanek M, Haluszko A, Mackiewicz M, Rapinski A, Leach S, Soriano I, Lillios KT, Cardoso JL, Pearson MP, Wlodarczak P, Price TD, Prieto P, Rey PJ, Risch R, Rojo Guerra MA, Schmitt A, Serralongue J, Silva AM, Smrcka V, Vergnaud L, Zilhao J, Caramelli D, Higham T, Thomas MG, Kennett DJ, Fokkens H, Heyd V, Sheridan A, Sjogren KG, Stockhammer PW, Krause J, Pinhasi R, Haak W, Barnes I, Lalueza-Fox C, Reich D. 2018. The beaker phenomenon and the genomic transformation of northwest europe. Nature 555: 190–196.

Patterson N, Moorjani P, Luo Y, Mallick S, Rohland N, Zhan Y, Genschoreck T, Webster T, Reich D. 2012. Ancient admixture in human history. Genetics 192: 1065–1093.

Pickrell JK, Pritchard JK. 2012. Inference of population splits and mixtures from genome-wide allele frequency data. PLoS Genet 8: e1002967.

Purcell S, Neale B, Todd-Brown K, Thomas L, Ferreira MA, Bender D, Maller J, Sklar P, de Bakker PI, Daly MJ, Sham PC. 2007. Plink: A tool set for whole-genome association and population-based linkage analyses. Am J Hum Genet 81: 559–575.

Reich D, Green RE, Kircher M, Krause J, Patterson N, Durand EY, Viola B, Briggs AW, Stenzel U, Johnson PL, Maricic T, Good JM, Marques-Bonet T, Alkan C, Fu Q, Mallick S, Li H, Meyer M, Eichler EE, Stoneking M, Richards M, Talamo S, Shunkov MV, Derevianko AP, Hublin JJ, Kelso J, Slatkin M, Paabo S. 2010. Genetic history of an archaic hominin group from denisova cave in siberia. Nature 468: 1053–1060.

Sagart L, Jacques G, Lai Y, Ryder RJ, Thouzeau V, Greenhill SJ, List JM. 2019. Dated language phylogenies shed light on the ancestry of sino-tibetan. Proc Natl Acad Sci U S A 116: 10317–10322.

Sikora M, Pitulko VV, Sousa VC, Allentoft ME, Vinner L, Rasmussen S, Margaryan A, de Barros Damgaard P, de la Fuente C, Renaud G, Yang MA, Fu Q, Dupanloup I, Giampoudakis K, Nogues-Bravo D, Rahbek C, Kroonen G, Peyrot M, McColl H, Vasilyev SV, Veselovskaya E, Gerasimova M, Pavlova EY, Chasnyk VG, Nikolskiy PA, Gromov AV, Khartanovich VI, Moiseyev V, Grebenyuk PS, Fedorchenko AY, Lebedintsev AI, Slobodin SB, Malyarchuk BA, Martiniano R, Meldgaard M, Arppe L, Palo JU, Sundell T, Mannermaa K, Putkonen M, Alexandersen V, Primeau C, Baimukhanov N, Malhi RS, Sjogren KG, Kristiansen K, Wessman A, Sajantila A, Lahr MM, Durbin R, Nielsen R, Meltzer DJ, Excoffier L, Willerslev E. 2019. The population history of northeastern siberia since the pleistocene. Nature 570: 182–188.

Siska V, Jones ER, Jeon S, Bhak Y, Kim HM, Cho YS, Kim H, Lee K, Veselovskaya E, Balueva T, Gallego-Llorente M, Hofreiter M, Bradley DG, Eriksson A, Pinhasi R, Bhak J, Manica A. 2017. Genome-wide data from two early neolithic east asian individuals dating to 7700 years ago. Sci Adv 3: e1601877.

Sun J, Zhao J, Cheng H-Z, Yang X, He G, Guo J, Li Y, Hu R, Chen G, Wei L-H, Wang C-C. 2020. Genetic affinity and substructure of hmong-mien speaking populations inferred from genome-wide array genotyping of three unrecognized ethnic groups dongjia, xijia and gejia. Mol Genet Genomics.

Wang C-C, Yeh H-Y, Popov AN, Zhang H-Q, Matsumura H, Sirak K, Cheronet O, Kovalev A, Rohland N, Kim AM, Bernardos R, Tumen D, Zhao J, Liu Y-C, Liu J-Y, Mah M, Mallick S, Wang K, Zhang Z, Adamski N, Broomandkhoshbacht N, Callan K, Culleton BJ, Eccles L, Lawson AM, Michel M, Oppenheimer J, Stewardson K, Wen S, Yan S, Zalzala F, Chuang R, Huang C-J, Shiung C-C, Nikitin YG, Tabarev AV, Tishkin AA, Lin S, Sun Z-Y, Wu X-M, Yang T-L, Hu X, Chen L, Du H, Bayarsaikhan J, Mijiddorj E, Erdenebaatar D, Iderkhangai T-O, Myagmar E, Kanzawa-Kiriyama H, Nishino M, Shinoda K-i, Shubina OA, Guo J, Deng Q, Kang L, Li D, Li D, Lin R, Cai W, Shrestha R, Wang L-X, Wei L, Xie G, Yao H, Zhang M, He G, Yang X, Hu R, Robbeets M, Schiffels S, Kennett DJ, Jin L, Li H, Krause J, Pinhasi R, Reich D. 2020a. The genomic formation of human populations in east asia. bioRxiv: 2020.2003.2025.004606.

Wang M, Zou X, Ye H-Y, Wang Z, Liu Y, Liu J, Wang F, Yao H, Chen P, Tao R, Wang S, Wei L-H, Tang R, Wang C-C, He G. 2020b. Peopling of tibet plateau and multiple waves of admixture of tibetans inferred from both modern and ancient genome-wide data. bioRxiv.

Wang M, Zou X, Ye H-Y, Wang Z, Liu Y, Liu J, Wang F, Yao H, Chen P, Tao R, Wang S, Wei L-H, Tang R, Wang C-C, He G. 2020c. Peopling of tibet plateau and multiple waves of admixture of tibetans inferred from both modern and ancient genome-wide data. bioRxiv: 2020.2007.2003.185884.

Weir BS, Cockerham CC. 1984. Estimating f-statistics for the analysis of population structure. Evolution 38: 1358–1370.

Yang MA, Fan X, Sun B, Chen C, Lang J, Ko YC, Tsang CH, Chiu H, Wang T, Bao Q, Wu X, Hajdinjak M, Ko AM, Ding M, Cao P, Yang R, Liu F, Nickel B, Dai Q, Feng X, Zhang L, Sun C, Ning C, Zeng W, Zhao Y, Zhang M, Gao X, Cui Y, Reich D, Stoneking M, Fu Q. 2020. Ancient DNA indicates human population shifts and admixture in northern and southern china. Science 369: 282–288.

Zhang M, Fu Q. 2020. Human evolutionary history in eastern eurasia using insights from ancient DNA. Curr Opin Genet Dev 62: 78–84.

Zhang M, Yan S, Pan W, Jin L. 2019. Phylogenetic evidence for sino-tibetan origin in northern china in the late neolithic. Nature 569: 112–115.

Zou X, Wang Z, He G, Wang M, Su Y, Liu J, Chen P, Wang S, Gao B, Li Z, Hou Y. 2018. Population genetic diversity and phylogenetic characteristics for high-altitude adaptive kham tibetan revealed by dnatyper(tm) 19 amplification system. Front Genet 9: 630.

